# Effect of anthropogenic vibratory noise on plant development and herbivory

**DOI:** 10.1101/2021.04.28.441746

**Authors:** Estefania Velilla, Laura Bellato, Eleanor Collinson, Wouter Halfwerk

## Abstract

A growth in anthropogenic activities and infrastructure has led to increasing subterranean vibratory noise levels. Inland wind energy turbines, which are mostly located in agricultural fields, are a fast growing source of vibrational noise. Plants, which are rooted in the soil are constantly exposed to windmill-induced vibrations propagating through the ground. We have little understanding on how anthropogenic seismic vibrations affect plant development and how that in turn can affect plant-insect interactions. In this study we investigated the effect of windmill-like underground vibrational noise on plant development and on a plant-herbivore interaction. We experimentally exposed *Pisum sativum* plants from seed stage to seed production stage to high and low vibrational noise levels and monitored them daily. We recorded germination, flowering and fruiting time, as well as daily shoot-length growth. Moreover, we tested the direct and indirect effects of vibrational noise on herbivory intensity by the generalist caterpillar *Spodoptera exigua*. We found that plants exposed to high vibrational noise grew significantly faster and taller than plants exposed to low vibrational noise. Additionally, plants treated with high noise germinated, flowered and produced fruits quicker than those treated with low noise. However, the differences in germination time, flowering time and fruiting time between the treatments were not statistically significant. Furthermore, we did not find an effect of vibrational noise on herbivory intensity. Vibrational noise could have consequences for both natural plant communities and agricultural crops by altering interspecific competition and by shifting growth-defence activation trade-offs.

## Introduction

The global increase in human infrastructure is well-known to affect ecosystems and the organisms that live in them. An important component of human activities associated with infrastructure is the production of substrate-borne vibrations. Road and railway traffic as well as construction work has led to an increase in subterranean vibratory noise levels, potentially influencing organisms that live in the soil. Organisms ranging from animals (Cocroft and Rodriguez 2005; Cocroft et al. 2014) to plants (Takahashi et al. 1991; Uchida and Yamamoto 2002; Gagliano et al. 2012, 2017; Ghosh et al. 2016) to fungi (Leach, 1980) are known to rely on vibratory signals and cues for their survival and reproduction (De Luca and Vallejo-Marín, 2013; Appel and Cocroft 2014; Veits et al. 2019). Human-induced vibratory noise levels may therefore impact organisms across trophic levels. Despite the far-reaching ecological implications, only a handful of studies have looked into the effect of vibrational noise and have mostly focused on animals (Wu and Elias 2014; Gagliano et al. 2017; Roberts and Elliott 2017; Caorsi et al. 2019).

The few studies on the effects of vibratory noise on plants suggest that vibrational cues are used across a range of different ecological contexts. *Zea mays* roots for example grow towards vibrations associated with flowing water (Gagliano et al. 2017) and *Arabidopsis thaliana* increase their anti-herbivory chemical defense when exposed to the chewing vibrations induced by their herbivores (Appel and Cocroft 2014). Moreover, stimulating plants with sounds and vibrations of different frequency ranges has been shown to affect germination timing (Uchida and Yamamoto 2002; Creath and Schwartz 2004; Cai et al. 2014) and growth (Takahashi et al. 1991). Besides the direct effects of vibrations on plants, vibratory noise can also affect organisms that interact with plants, hence indirectly affecting plants via species interactions. For example, vibrations may deter beneficial soil fauna such as earthworms and therefore decrease soil fertility, or it can affect organisms in the trophic cascade of herbivores, predators and parasitoids.

Wind energy turbines are a fast-growing potential source of anthropogenic vibrational noise. In Europe, wind energy remains the second largest form of power generation capacity, with a total net installed capacity of 168.7 GW (Wind in power 2017 report). Most inland wind energy turbines are located in agricultural fields where plants and the insects that interact with them are potentially constantly exposed to vibrational noise.

In our current study we assessed whether vibrational noise generated by wind energy turbines influences plant-insect interactions. We first examined the effect of vibrational noise on plant developmental processes (e.g. germination, flowering and fruiting time, and growth). In the plant development experiment, we were interested to determine whether plants exposed to vibrational noise germinated, flowered and fruited sooner or later than plants exposed to low vibrational noise, and whether there were differences in growth between the plants exposed to high or low vibrational noise. Next, we tested for direct and indirect effects of vibrational noise on herbivory. We experimentally exposed *Pisum sativum* seeds to high and low vibrational noise and tracked their development on a daily basis. After fruiting of the plants, we carried out an herbivory experiment, using the generalist caterpillar *Spodoptera exigua* and full-factorial design. For the herbivory experiment we were interested in two questions: 1) does vibrational noise occurring at the moment of foraging affect herbivory? and 2) Does prior plant exposure to vibrational noise have carry-over effects on herbivory? For example, via changes in secondary metabolites? Caterpillars were placed on plants that either developed during low or high noise conditions, and were exposed to either high or low vibrational noise during the foraging period.

## Materials and Methods

### Study species

We decided to work with *P. sativum* plants because we were interested in linking our vibrational noise field measurements to crop plants and because of evidence indicating the sensitivity of this species to vibrational noise (Gagliano et al. 2017). Organic *P. sativum* seeds were obtained from the company Intratuin (Nobelweg 10, 1097 AR Amsterdam, The Netherlands).

For our herbivory experiments we used the generalist caterpillar *S. exigua*. We obtained *S. exigua* eggs from the company Entocare (Haagsteeg 4, 6708 PM Wageningen, The Netherlands). The animals were reared in climate rooms at 26°C ± 1°C and 80% relative humidity on a 12D/12N light cycle. Caterpillars were fed ad-libitum with a corn-based artificial diet. For our herbivory experiments we used caterpillars in the 5^th^ instar. Caterpillars that took part in our experiments were fed with *P. sativum* the day before testing them to acclimatise them to the diet.

### Experimental procedures

We exposed *P. sativum* to high and low levels of vibratory noise throughout their development (47 ± 3.5 days). Seeds were planted individually on 26 June 2019 in 6 cm diameter plastic plots containing organic potting soil (Horticoop, Kalppolder 150, Bleiswijk, The Netherlands). The entirety of this experiment took place in a climate chamber at 20°C with an 8:16 (L:D) photoperiod and 70% relative humidity. The lighting in the experimental room contained Photo Active Radiation (PAR) in the range of 400-700 nm and some far-red lighting using LED lamps, in the range of 600-800 nm. The light level in the experimental room was 450 µmol/m^2^ /s. We exposed a total of 40 plants to vibrational noise from seed stage to fruiting stage. From the 40 plants, 20 were exposed to high vibrational noise and the other 20 were exposed to low vibrational noise (see below). Ten pots were placed on a plastic tray (48.5 x 31cm). Below the tray, four shakers (Monacor AR-30, Monacor International GmbH & Co. Zum Falsch 36, 28307 Bremen, Germany) connected to a constantly charging mp3 player (Lenco Xemio-200) and to a stereo amplifier (Renkforce SAP-702 2 x 20 W manufacturer No. RF-3511635 or Renkforce T21 2 x 50 W manufacturer No. RF-4602693) broadcasted vibrational pink noise.

Our playback stimulus (pink noise) was based on field recordings of subterranean vibrational noise induced by windmills (see Velilla *et al* 2021, for methods on field recordings of wind turbine-induced vibrations). Since windmill-induced vibrational noise was highly represented by energy in the low frequencies (< 100Hz), we decided to use pink noise as a standardized playback stimulus, also biased towards the low frequencies, albeit in a wider range.

We measured the ambient vibratory noise levels of our setup using a Laser-Doppler vibrometer (LDV; Polytec PDV-100, set to 100 mm/s/V, sampling rate 22kHz) connected to an oscilloscope (Rigol, DS1054, 4 Channel 50MHz, 1GSa/s). We adjusted the amplitude by changing the settings of the amplifier. For the low-noise treatment we matched the RMS values obtained from the oscilloscope to the value obtained for the ambient conditions of the room. For the high noise treatment, we set the amplitude 12 dB higher compared to the low noise treatment. A difference of 12 dB corresponds to the difference obtained from recording windmill induced soil vibrations at 8 m compared to 128 m. We chose 12 dB since this was the maximum intensity-difference we could obtain with our speaker setup without creating distortion. The amplitude of our vibrational noise stimuli was calibrated on several positions on the tray to ensure that all plants were exposed to equal vibrational noise amplitude (Fig. 1).

**Fig. 1.**
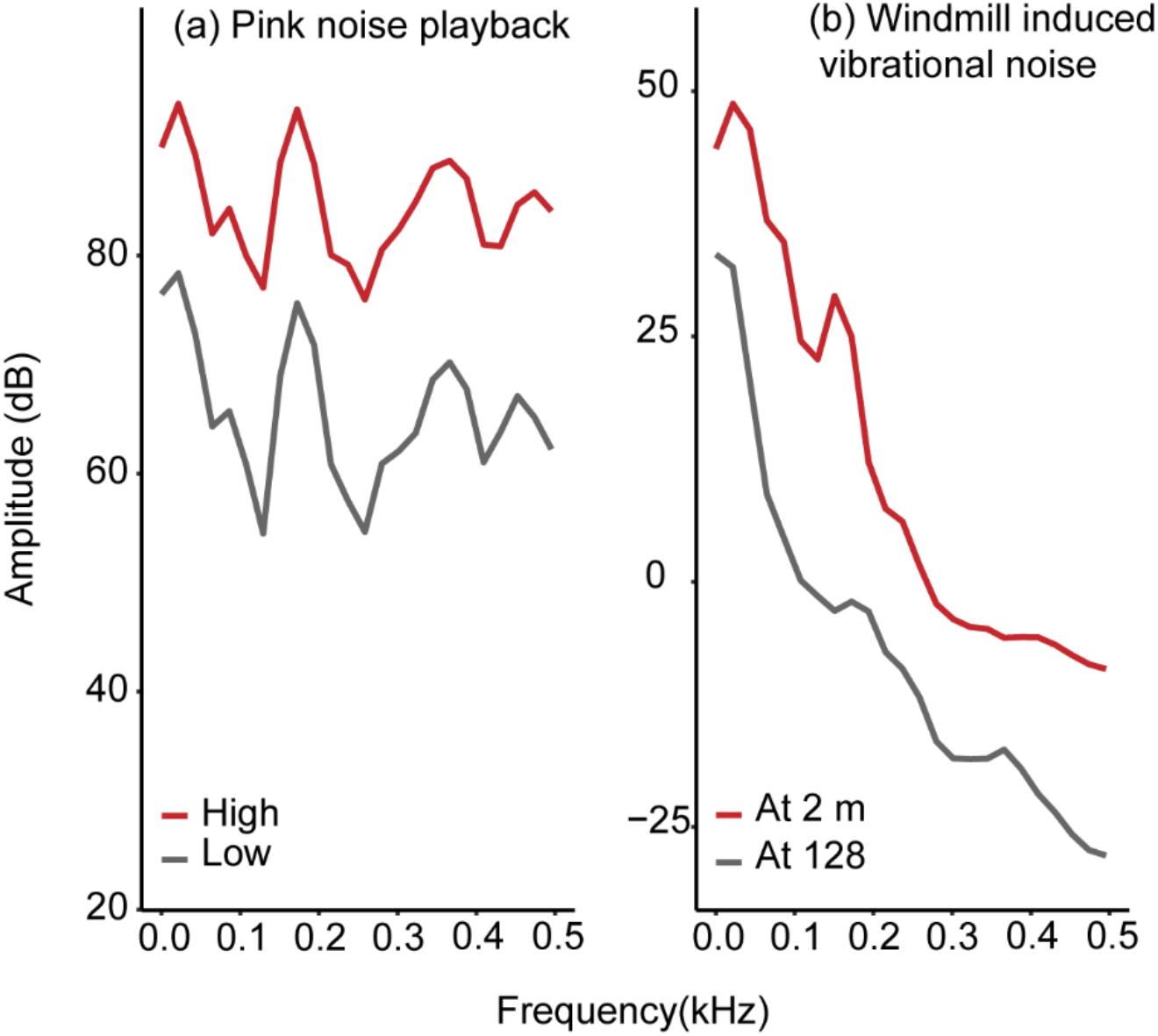
Power spectra of experimental playback stimulus and field recordings. **(a)** Power spectra from recordings of vibrational pink noise playback stimulus (“Hanning” window, window size = 2048) recorded with a Doppler laser vibrometer on the centre of an experimental tray containing 10 potted *P. sativum* plants. The high noise treatment (red line) was 12dB louder than the low noise treatment, and the low noise treatment (dark grey line) was approximately 2dB louder than ambient silent conditions calculated from root mean square readings from an oscilloscope. **(b)** Power spectra from normalized recordings (“Hanning” window, window size = 2048) of windmill-induced vibrational noise in the field at 2m (red line) and 128m (dark grey line) from the base of the wind turbine. Recordings were made with a vertical geophone.

**Fig. 1.**
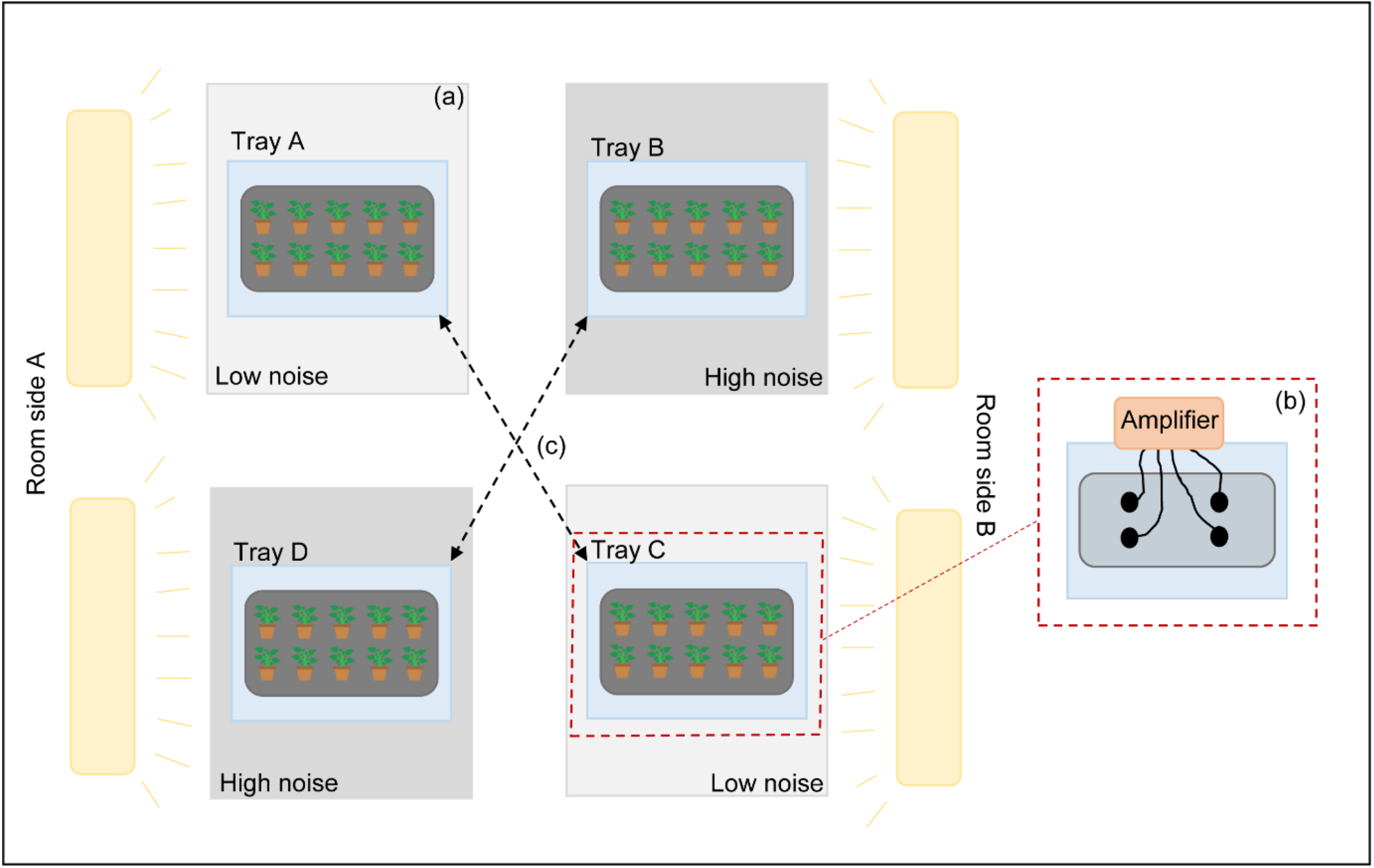
Graphical depiction of experimental set-up where we tested the effect of high and low vibratory noise on plant development. (a) Experimental table (light grey) with a plastic tray (blue) with four shakers on it. On top of the shakers another tray (dark grey) contained 10 *P. sativum* potted seeds/plants. We had two tables per treatment placed opposite and diagonally to each other. (b) the four shakers were attached to an amplifier, and the experimental stimulus was played from a constantly charging mp3 player attached to the amplifier. (c) We switched the position of the trays (light blue) 19 days after planting to control for confounding effects of room side.

We divided the treatments in two trays and the trays were positioned opposite to each other (on opposite sides of the room), to control for any lighting effects (Fig. 2). Halfway through the experiment (19 days after planting) we switched tray position so that plants in one side of the room would be exposed to the other side of the room, this way avoiding confounding effects of location in the room.

The seeds/plants were checked on a daily basis except for weekends. We extrapolated based on developmental events recorded during the working week to calculate the day an event took place during the weekend. We noted germination date, date of appearance of first flower and date of appearance of first fruit. Additionally, shoot length was measured on a daily basis with a metallic 30 cm-ruler. Plants were watered by adding water to their tray allowing them to absorb as much water as needed.

For the herbivory experiment we exposed caterpillars to high and low vibrational noise while they foraged on plants that had previously been exposed to either high or low vibrational noise during the plant development experiment (Fig. 3). Half of the plants that were exposed to high vibrational noise during the plant development experiment were exposed to low vibrational noise during the herbivory experiment, while the other half were exposed to high vibrational noise. The same applied for plants exposed to low vibrational noise during the plant development experiment, with half of the plants exposed to high vibrational noise during the herbivory experiment, and the other half exposed to low vibrational noise. We used the change in caterpillar weight (delta weight) as a proxy to herbivory intensity. Some caterpillars failed to forage at all and were excluded from the results. The caterpillars that failed to forage fell into the soil and did not climb the plant again. The number of caterpillars that failed to forage was distributed across treatments. In total we obtained data for 35 caterpillar-plant combinations.

**Fig. 3.**
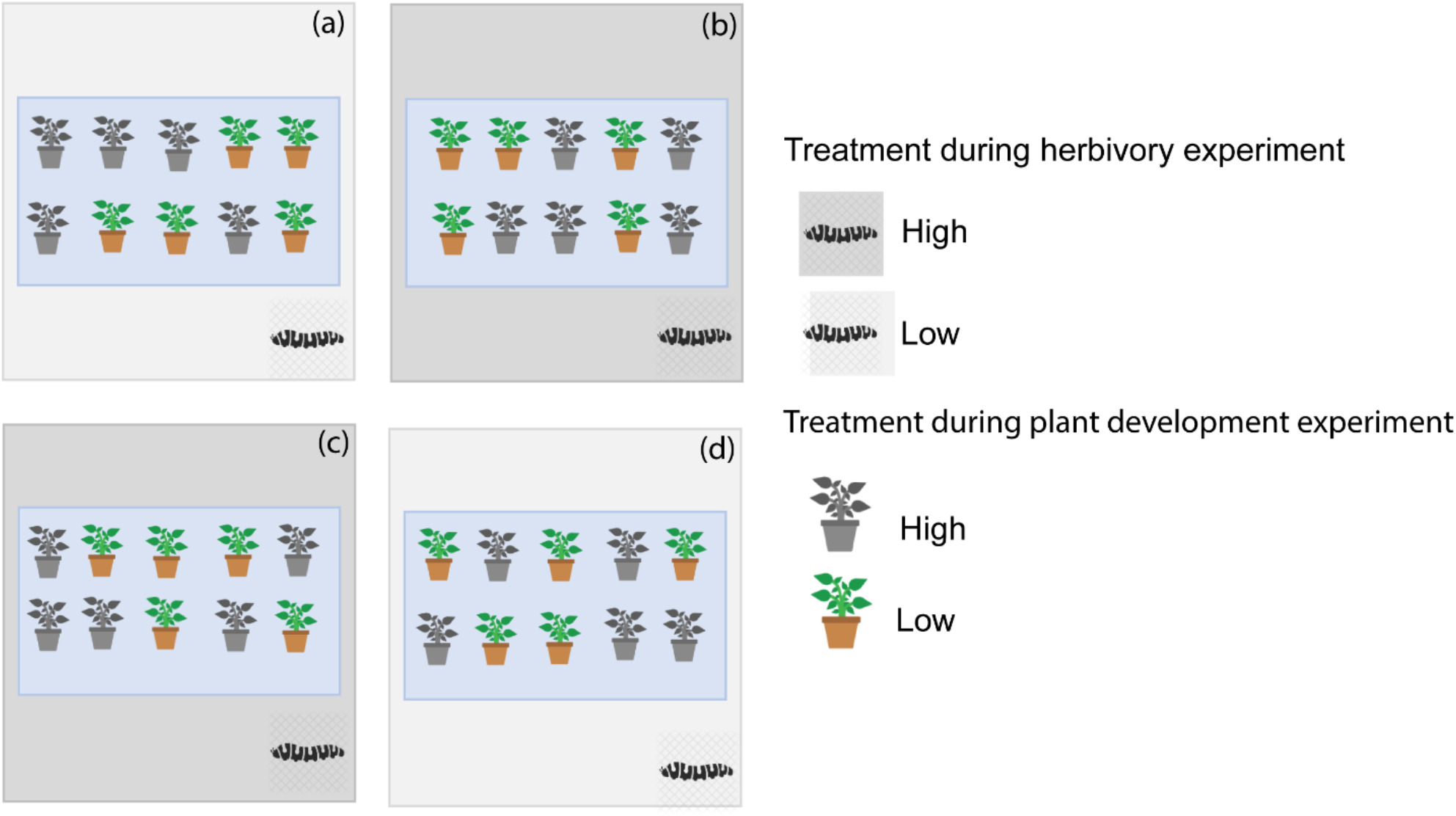
Graphical illustration of herbivory experiment setup. We tested the effect of vibratory noise on herbivory of P. sativum by S. exigua. Noise could affect foraging in caterpillars directly, or indirectly via effects on plants previously exposed noise. Therefore, caterpillars were tested on the plants used during the plant development under noise conditions experiment. Caterpillars were placed on the leaves of plants and exposed to high or low noise conditions for 24 hours. (a) Half of the plants on this tray had been exposed to high noise conditions during the plant developing experiment, and the other half to low noise conditions. (b) Half of the plants on this tray had been exposed to high noise conditions during the plant developing experiment, and the other half to low noise conditions. (c) and (d) are replicates of (a) and (b), and therefore the same description applies. Plants were randomly (balanced) placed on the experimental trays.

We deprived the caterpillars (5^th^ instar, 22-27 days old) of food for 2 hours before testing them. Caterpillars were weighed just before placing them on a plant. Individual shakers were placed under individual plants. Caterpillars were allowed to forage for 24 hours, after which they were removed and immediately weighed. Foraging intensity by caterpillars under different noise levels could differ because caterpillars might be disturbed by noise, in which case we would expect foraging intensity to be lower under high noise conditions, with caterpillars showing a lower weight increase. Alternatively, caterpillar chewing vibrations which can increase secondary metabolite emission in plants (Appel and Cocroft 2014) could be masked by our high noise treatment. The masking of caterpillar-induced vibrations could result in lower emission of secondary metabolites, thus allowing caterpillars to forage for a longer amount of time. In this case, we would expect a higher caterpillar weight increase. For logistical reasons we did not measure surface eaten from the leaves. However, we expect surface eaten to be highly correlated with increases in caterpillar weight. Therefore, we used delta weight as a proxy to herbivory intensity.

### Data analyses

Statistical analyses were done with R version 3.3.1 (R Core Team, 2016), run in the R studio interface (RStudio Team, 2015).

We used a Kruskal Wallis test to test the effect of high and low vibrational noise on the number of days it took for seeds to germinate. We used a non-parametric test here because our data did not follow normal distribution, and no transformation or alternative choice of distribution family produced a satisfactory model. Furthermore, to test the effect of high and low vibrational noise on the number of days it took for to produce their first flower and their first fruit we used linear mixed effects models following Gaussian distribution. We included tray ID as a random effect with a random intercept. Days to flowering and days to fruiting, which were our response variables were both squared-root transformed to achieve a better model fit. The number of days it took to flower and produce fruit was calculated from germination date and not from planting date. To test the effect of high and low vibrational noise on plant growth we used a linear mixed effects model with repeated shoot length measurements as our response variable. We nested plant identity in tray as a random effect, with a random intercept.

We tested whether vibrational noise affected herbivory using a linear regression, with caterpillar delta weight as our response variable. Our main predictors were treatment during the herbivory experiment (high or low vibrational noise at the time of foraging), treatment during the plant development experiment (treatment to which the plants had been exposed to in the previous experiment) and final shoot length. We tested the interaction between treatment and previous treatment. We included plant final shoot length as a predictor to control for effects of plant size on the investment of plant defences against herbivory.

All model residuals were inspected by means of Q-Q plots and histograms. There were no deviations from the normality or variance homogeneity assumptions from linear models. The effect of individual model predictors was established by full-null model comparisons by means of Wald Chi-square statistics from the “Anova” function in the statistical package “Car” (Fox et al., 2012).

## Results

### Vibrational noise affects plant growth and flowering time

Our repeated measures analysis using shoot length as a proxy for growth revealed that vibrational noise had a significant effect on growth, with plants growing taller under high vibratory noise conditions (LMM, high vibrational noise treatment, n = 39, *β* = 2.9525, *t =* −2.191, *p* = 0.03; Fig. 4).

**Fig. 4.**
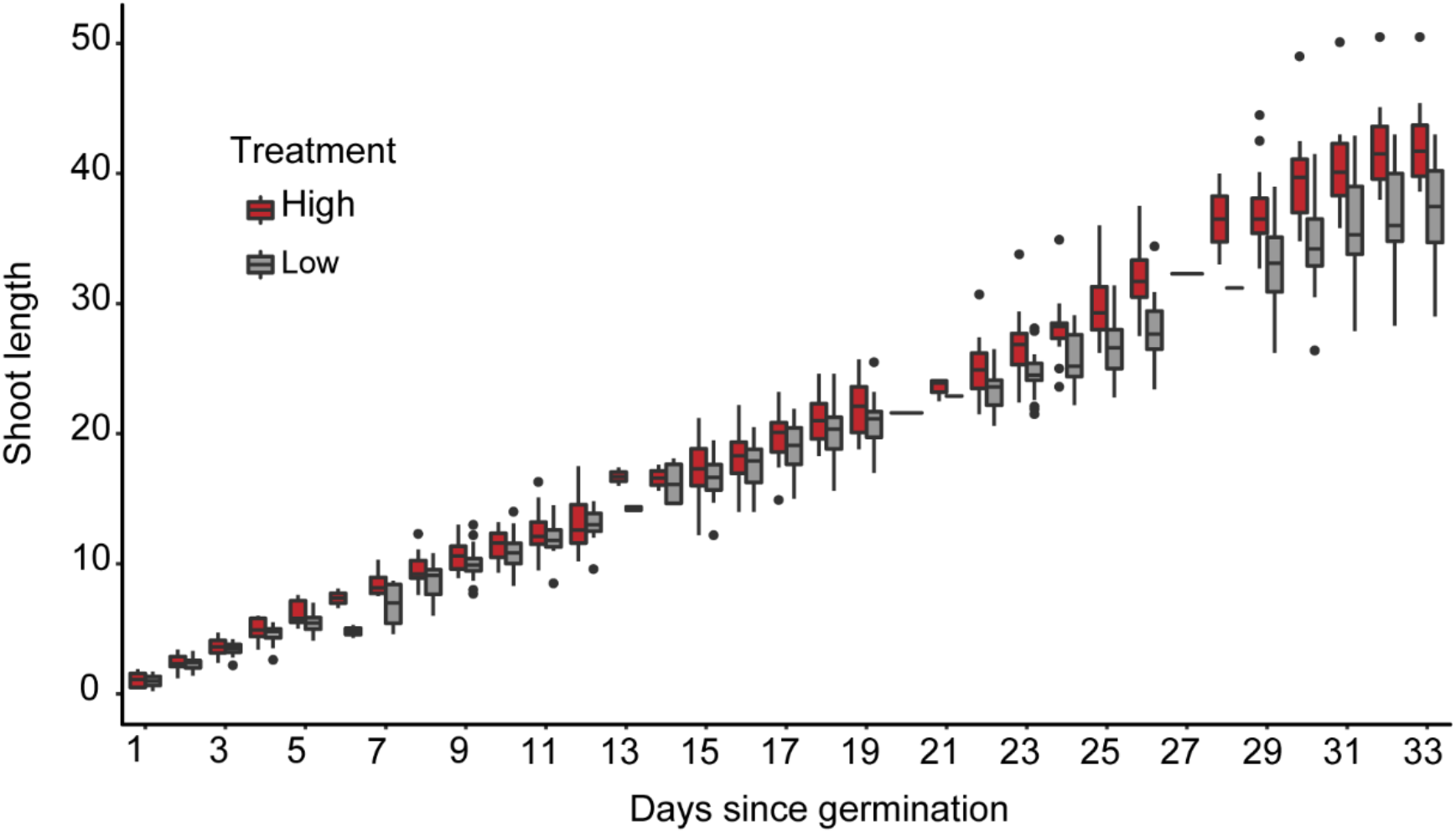
Boxplot showing increase in shoot-length (cm) over time (days since germination) for plants growing under high (red) and low (grey) vibrational noise. Plants exposed to high (red) vibrational noise grew taller than plants exposed to low (grey) vibrational noise. The interquartile range was taken as the range from 0-25th percentile. From the mean, the whiskers show the highest and lowest value within 1.5 times the interquartile range.

Although seeds exposed to high noise germinated slightly earlier than seeds exposed to low noise, these differences were not statistically significant (n= 39, Kruskal Wallis, χ^2^_(1)_ = 0.11, *p*= 0.73, Fig. 5 (a)). Noise treatment also tended to affect flowering time (LMM, *df =* 1, *F=* 3.24, *p =* 0.08), with plants exposed to high noise conditions flowering on average around 2 days earlier than plants exposed to low noise conditions. Plants treated with high vibratory noise produced their first flower on average within 37.2 (SD ± 4.2) days after germination, while the mean number of days it took plants exposed to low vibratory noise to produce their first flower was 39.7 (SD ± 4.2) days after germination (Fig. 5 (b)). Fruiting time did not differ significantly between the treatments (LMM, *df* = 1, *F =* 2.0476, *p* = 0.16, Fig. 5 (c)).

**Fig. 5.**
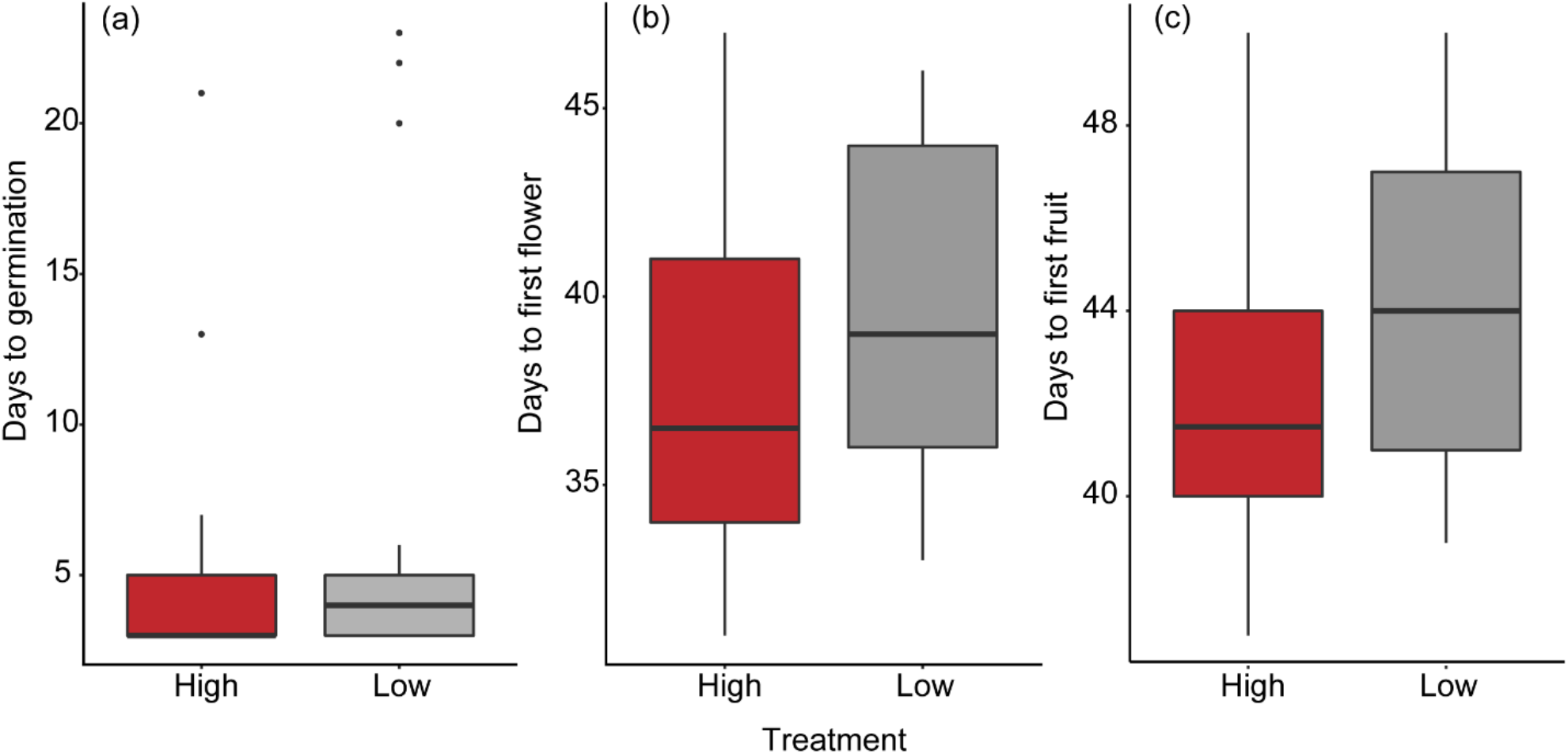
(a) Boxplot showing the number of days it took seeds exposed to high vs. low vibratory noise to germinate. Seeds exposed to high vibratory noise germinated approximately 1 day earlier than seeds exposed to low vibratory noise. (b) Number of days it took plants exposed to high vs. low vibratory noise to produce their first flower. Plants exposed to high vibratory noise produced their first flower approximately 2 days earlier than seeds exposed to low vibratory noise. The difference was near-significant. (c) Number of days it took plants exposed to high vs. low vibratory noise to produce their first fruit. Plants exposed to high vibratory noise produced their first flower approximately 1.5 days earlier than seeds exposed to low vibratory noise. The difference was not significant. The difference was not significant. The interquartile range was taken as the range from 0-25th percentile. From the mean, the whiskers show the highest and lowest value within 1.5 times the interquartile range.

### Vibrational noise does not affect herbivory

We tested whether vibrational noise affected herbivory directly or indirectly. We found no differences in herbivory intensity between caterpillars foraging in high versus low vibrational noise. Furthermore, we examined whether the vibrational noise treatment during the plant growth experiment affected herbivory intensity. We found no carry over effects (Linear regression, *df=* 1, treatment during herbivory experiment, *F =* 0.0263, *P =* 0.87; treatment during plant development experiment, *F* = 0.7338, *P =* 0.39; interaction between treatment during herbivory experiment and treatment during plant development experiment, *F* = 0.3648, *P* = 0.55; shoot length, *F=* 0.0183, *P =* 0.89, Fig. 6).

**Fig. 6.**
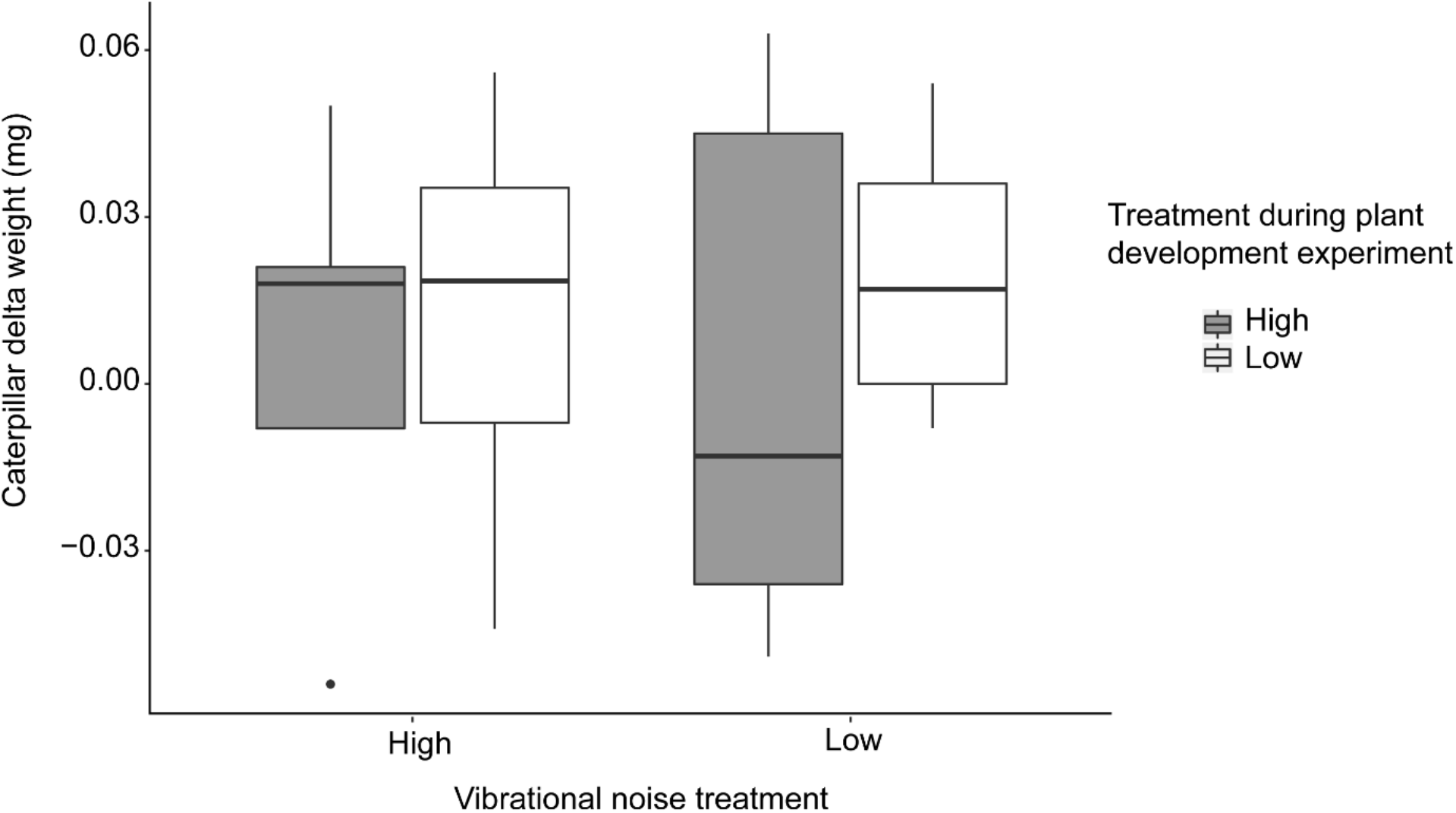
Experiment testing the effect of noise on herbivory intensity of *S. exigua* caterpillars on *P. sativum* plants. This boxplot shows changes in caterpillar weight after foraging under high or low vibratory noise. The plants used in this experiment were the same plants used in the experiment that tested the effect of vibratory noise on plant development. To disentangles whether the effect on herbivory was a direct effect on the caterpillars or a carry-over effect from the plants from noise exposure during the previous experiment, half of the plants that were exposed to high noise during the plant development experiment were exposed to low noise during the herbivory experiment, and the other half to high noise. In the same manner, half of the plants that were exposed to low noise during the plant development experiment were exposed to high noise during the herbivory experiment, and the other half to low noise (Fig. 3). The difference was not significant. The interquartile range was taken as the range from 0-25th percentile. From the mean, the whiskers show the highest and lowest value within 1.5 times the interquartile range.

## Discussion

The aim of this study was to examine whether windmill induced vibrations can influence plants and their herbivores. In a first experiment we tested whether vibrational noise affected plant developmental processes like germination, flowering time, fruiting time, and growth. Using the same plants, we tested in a follow-up experiment whether vibrational noise affected herbivory.

Our results show that plant development can be affected by vibrational noise that is reminiscent of noise profiles found in the proximity of wind power turbines. Plants exposed to high noise levels grew faster and taller compared to plants exposed to low noise levels. Furthermore, we found that on average plants exposed to high noise germinated and flowered slightly faster than plants exposed to low noise. However, the differences between treatments were not statistically significant.

Although we did not find statistical differences in germination, studies on other plants have repeatedly shown that vibrations can influence this important developmental stage. Mechanical stimulation with 50Hz vibrations has e.g. been shown to promote germination in *Cucumis sativus* and *Oryza sativa* (Takahashi et al. 1991). Uchida and Yamamoto (2002) also showed that vibrational stimulation with monotone frequencies in the range of 40-120 Hz affected germination of *Arabidopsis thaliana.* Ethylene synthesis, which is necessary for seed germination (Abeles et al. 1992), is known to be induced by mechanical stresses like bending, rubbing and shaking of plants (Goeschl et al. 1966; Takahashi and Jaffe 1984), and more recently shown to be induced by vibrations (Uchida and Yamamoto 2002). Some studies have also provided evidence that vibrations of certain frequencies positively influence not only germination, but also root elongation, callus growth, and cell cycling (Gagliano 2013; Chowdhury et al. 2014; Teixeira da Silva and Dobránszki 2014). It could be that *P. sativum* is sensitive to very low frequency vibrations (< 50Hz), as was shown for *C. sativus* and *O. sativa* (Takahashi et al. 1991). Given that the frequency response of our shakers was poor for the low frequencies (< 100Hz), it is possible that the effect of our treatments was hindered by limitations in our playback setup. Alternatively, it is possible that our sample size was too small to detect an effect of our treatment on germination, flowering and fruiting time. We would not have experienced the same problem when we tested daily growth, because we used a repeated measures analysis, which typically have higher statistical power (Guo et al. 2013).

In our study we show that plants exposed to the high vibrational noise treatment experienced significantly higher growth than plants exposed to low vibrational noise. These different developmental trajectories may have already been initiated at the start of germination. Our results seem therefore consistent with previous studies, which have shown that vibrations stimulate the growth and development of several plant species (Takahashi et al. 1991; Collins and Foreman 2001; Mishra et al. 2016).

Vibrations are not only important for plant developmental processes, but they can also be important for plant-animal interactions. For instance, *Arabidopsis thaliana* increases its anti-herbivory defences when stimulated with their herbivores’ chewing vibrations (Appel and Cocroft 2014), and *Oenothera drummondii* increases its nectar sugar concentration when exposed to the buzzing vibrations of its pollinator (Veits et al. 2019). We wanted to investigate whether vibrational noise affected plant-insect interactions, for interactions in which vibrations provided valuable information. Plants that are constantly exposed to vibrational noise, like plants in a windmill field, could have a harder time detecting the chewing vibrations of their herbivores making them more vulnerable to higher herbivory intensity. In this study, however, we found that vibrational noise did not affect herbivory intensity and we did not see any evidence of carry over effects on herbivory from previously exposing plants to high or low vibrational noise. Although we did not see a herbivory effect in our experiments, it is important to point out that we used caterpillar delta weight as a proxy to herbivory intensity, which might not be the most accurate measurement of effects on herbivory, since fluctuations in weight might also be related to the individual moving activity patterns of the caterpillars.

In conclusion, we found that *P. sativum* germinates, flowers and produces fruit slightly quicker when stimulated with higher amplitude non-monotonic low frequency vibrational noise than when stimulated with a lower amplitude of the same noise stimulus. Furthermore, we found that *P. sativum* plants exposed to high vibrational noise grew significantly taller than plants exposed to low vibrational noise. The amplitude difference at which we see this effect is ecologically relevant for plants growing in fields where windmills are located.

The effect of sound vibrations on plants can be frequency and amplitude dependent (Chowdhury et al. 2014). Consequently, not all plant species’ development could be affected by windmill-induced vibrations in the same way, possibly altering competition rates between plant species. Although this would not necessarily be a problem for cultivated fields with monocultures, it could have consequences for natural plant communities where interspecific competition is an important determinant of the structure and the dynamics of plant communities (Aerts 1999). Furthermore, investment in plant growth might come at the expense of plant defence activation, which is imperative for plant survival (Huot et al. 2014), since plants count with a limited pool of resources that can be used for growth or for defence (Coley et al. 1985; Simms and Rauscher 1987; Herms and Mattson 1992). Our results highlight the susceptibility of plants to vibratory noise. Future studies are needed to understand the sensitivity of plants to vibrations of various frequencies, and whether different developmental stages are affected differently by the different frequencies.

Finally, the positive effects on plant growth reported in this study may offset some of the potentially negative effects we observed at the field sites from which we obtained the seismic noise recordings. At these organic crop fields, we observed earth worm abundances to decrease with increasing seismic noise levels (Velilla *et al.* 2021). As earthworms are well known ecosystem engineers influencing soil functioning, a negative effect on their populations may reverberate into reduced crop performances. Future studies should combine both direct and indirect effects on plants, animals and perhaps even micro-organisms to get a better understanding of the impact of seismic noise on soil ecosystems.

## Acknowledgements

We thank Richard van Logtestijn for his help setting up the plant climate chambers and Hans Cornelissen for his advice on plant choice.

